# Impact of genetic susceptibility to multiple sclerosis on the T cell epigenome: proximal and distal effects

**DOI:** 10.1101/2020.07.11.198721

**Authors:** Tina Roostaei, Hans-Ulrich Klein, Daniel Felsky, Pia Kivisäkk, Sarah M. Connor, Alexandra Kroshilina, Christina Yung, Yiyi Ma, Belinda J. Kaskow, Xiaorong Shao, Brooke Rhead, Jia Liu, Nikolaos Patsopoulos, Lisa F. Barcellos, Howard L. Weiner, Philip L. De Jager

**Affiliations:** Center for Translational and Computational Neuroimmunology, Department of Neurology and the Taub Institute for Research on Alzheimer’s disease and the Aging brain, Columbia University Irving Medical Center; Krembil Centre for Neuroinformatics, Centre for Addiction and Mental Health, University of Toronto; Alzheimer’s Clinical and Translational Research Unit, Department of Neurology, Massachusetts General Hospital; Ann Romney Center for Neurologic Diseases, Brigham and Women’s Hospital, Harvard Medical School; Genetic Epidemiology and Genomics Laboratory, University of California, Berkeley; Advanced Science Research Center at the Graduate Center, Neuroscience Initiative, City University of New York

## Abstract

We establish a genome-wide map of DNA methylation quantitative trait locus (mQTL) effects in CD4^+^ T cells isolated from multiple sclerosis (MS) patients. Utilizing this map in a colocalization analysis, we identify 19 loci where the same haplotype drives both MS susceptibility and local (*cis*-) DNA methylation. We also identify two distant (*trans*-) mQTL effects of MS susceptibility loci: (1) a chromosome 16 MS locus affects *PRDM8* methylation (a chromosome 4 region not previously associated with MS susceptibility), and (2) the aggregate effect of MS variants in the major histocompatibility complex (MHC, chromosome 6) influences DNA methylation near *PRKCA* on chromosome 17. Both effects are replicated in independent samples. Overall, we present a new methylome-wide mQTL resource for a key cell type in inflammatory disease research, uncover functional consequences of MS susceptibility variants, including the convergence of MHC risk alleles onto a new gene target involved in predisposition to MS.

## INTRODUCTION

Multiple sclerosis (MS) is a genetically complex inflammatory disease of the central nervous system. Despite a growing list of drugs that prevent relapses^1^, there is, as yet, no preventive strategy for MS, highlighting our limited understanding of the molecular events leading to disease onset. Given that genetic risk factors can be presumed to be causally linked to MS, they serve as a robust starting point for understanding the mechanisms predisposing to the disease. The most recent genome-wide association study (GWAS) of MS susceptibility^2^, provides compelling evidence for the effects of 32 independent susceptibility variants in the major histocompatibility complex (MHC) and 201 non-MHC susceptibility variants.

Our understanding of the functional consequences of these variants remains limited^2–5^. Here, we examined the first level of downstream molecular changes: alterations in the epigenome. Specifically, we focused on mapping methylation of CpG dinucleotides, an epigenomic mark for which nucleotide-resolution data can be reliably produced throughout the genome. Insights into the functional consequences of risk alleles on the epigenome can, in turn, be used to further elucidate the causal chain of biological mechanisms involved in disease onset. While the effects of many variants are shared, others demonstrate cell-type and context specificity. As CD4^+^ T cells are believed to play a major role in the pathogenesis of MS and other inflammatory disorders^3^, we purified these cells from MS patients to study the epigenome in a disease-relevant context.

CD4^+^ T cells have been interrogated in a number of recent quantitative trait locus (QTL) studies, albeit mainly from a gene expression (eQTL) perspective and mostly in healthy controls^3,6^. Here, using the Illumina MethylationEPIC array, we generated genome-wide DNA methylation profiles from CD4^+^ T cells isolated from 156 MS patients. These data were used to generate a **Resource** outlining the genome-wide genetic architecture of T cell DNA methylation levels in a disease state. We then performed a comprehensive set of analyses to determine the effects of MS risk loci. As a result, we (1) provide a genome-wide *cis*-DNA methylation QTL (mQTL) map of CD4^+^ T cells in MS patients, which can be utilized in future studies of MS and other inflammatory diseases, (2) identify *cis*-effects of MS genetic susceptibility variants on nearby CpG dinucleotide methylation, (3) discover and validate a *trans*-mQTL effect of an MS variant, and (4) demonstrate that polygenic scores of MS susceptibility influence DNA methylation at specific CpG dinucleotides and suggest the convergence of the effects of multiple variants on methylation levels in distal CpG sites.

## RESULTS

### Data generation

We selected subjects from participants in the Comprehensive Longitudinal Investigation of Multiple Sclerosis at the Brigham and Women’s Hospital (CLIMB) study^7^ that fulfilled our selection criteria: (1) age 18-55 years old, (2) a diagnosis of MS fulfilling 2010 McDonald criteria, (3) a relapsing-remitting disease course at the time of sampling, (4) being on one of two disease-modifying therapies (either glatiramer acetate [GA] or dimethyl fumarate [DMF]) at the time of sampling, (5) no evidence of disease activity in the prior 6 months, (6) no steroid use in the preceding 30 days, and (7) an Expanded Disability Status Scale (EDSS) score between 0 to 4. A prospectively collected, cryopreserved vial of peripheral blood mononuclear cells (PBMC) was accessed for each patient, and CD4^+^ T cells were purified after thawing using a positive selection strategy and a magnetic bead-based approach (**Methods**). DNA methylation profiles were generated using the DNA extracted from the purified CD4^+^ T cells using the Illumina MethylationEPIC array. We performed the mapping of mQTL effects using data from the 156 subjects for which both genotype and DNA methylation data passed all quality control measures (see **Methods**; demographic details are available in **Supplementary Table 1)**.

### Genome-wide mapping of *cis*-mQTLs in primary CD4^+^ T cells of MS patients

From the 769,699 DNA methylation sites that passed quality control measures, we found evidence for the influence of *cis* genetic effects (within ±1Mb of each CpG site) on the methylation levels of 107,922 CpGs (FDR-adjusted *p* <0.05) (**Supplementary Table 2**), after adjusting for the effect of technical and confounding variables (**Methods**). The high percentage of mCpGs (CpG sites which methylation levels are influenced by mQTLs = 14%) is comparable to findings from previous mQTL studies^8,9^. As expected, the majority of top *cis*-mQTL SNPs (mSNPs) are in a few kilobase distance from their target CpGs^8^, with 50% of mSNPs in 8kb and 90% in 123kb of their respective CpG sites (**Supplementary Figure 1**). To examine enrichment of *cis*-mCpGs in specific functional regions of the genome, we performed enrichment analyses using chromatin state annotations modeled for CD4^+^ T cells (sample #E043, Roadmap Epigenomics Project^10^), in addition to annotations for CpG islands and gene transcribed regions. We found lower than expected number of mCpGs in transcription start sites (TSS), transcribed regions, and CpG islands (which often occur at or near TSS). In contrast, enrichment was observed for mCpGs in flanking areas to TSS, flanking areas to CpG islands (island shores), and enhancer regions (**Figure 1A**).

**Figure 1.**
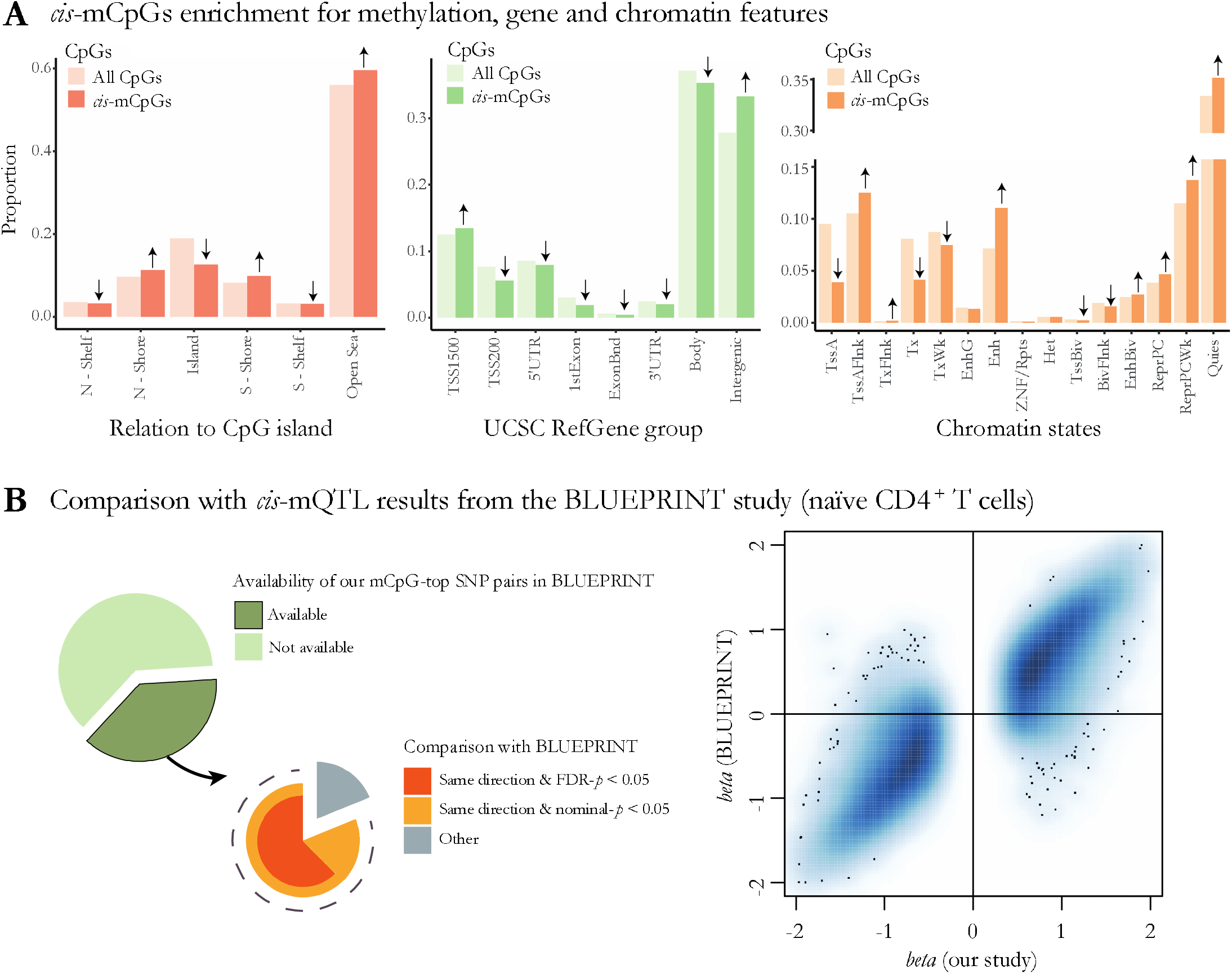
Characterization of genome-wide significant *cis*-mCpGs and *cis*-mQTLs. **(A)** Enrichment of the identified *cis*-mCpGs in comparison to all tested CpGs in relation to UCSC CpG islands and gene functional regions (annotations from R IlluminaHumanMethylationEPICanno.ilm10b2.hg19 package) and chromatin states modeled for CD4^+^ T cells (annotations from Roadmap Epigenomics Project, sample #E043). Significant enrichment/depletion are shown using upwards/downwards arrows, respectively. **(B)** Percentage of our identified *cis*-mCpGs available in the BLUEPRINT study and the comparison between the two studies regarding the direction of effect and significance of association for the top mSNPs are shown on the *left*. Comparison between the effect sizes found in our study and in BLUEPRINT are shown on the *right*.

We then compared our results with findings from a previous *cis*-mQTL study of naïve CD4^+^ T cells from 132 healthy controls performed as part of the BLUEPRINT Epigenome Project^9^. Of note, the Illumina 450K array used for measuring DNA methylation in the BLUEPRINT study contains only half of the number of CpGs tested in our study. Hence, our comparison is limited to the subset of CpGs shared between the two arrays. Of our 107,922 identified mCpG-mSNP pairs, 40,861 (38%) were tested in the BLUEPRPINT study. Of those, 38,782 (95%) showed the same direction of effect, 33,133 (81%) also had a nominally significant *p*-value (*p* <0.05), and 25,548 (63%) had an FDR-adjusted *p* <0.05 in the BLUEPRINT data (**Figure 1B**). The agreement between the two studies is further visualized using the estimated standardized regression coefficients for the effects of SNPs on the methylation of target CpGs (**Figure 1B**). The slight differences between our results and the BLUEPRINT study might have arisen from technical variation and population-specific effects, as well as the choice of cell type (CD4^+^ T cells comprising both naïve and memory cells in our sample vs. naïve-only CD4^+^ T cells in the BLUEPRPINT study) and disease status (MS patients in ours vs. healthy controls in the BLUEPRINT study). Nonetheless, the high degree of agreement between the two studies provides validation for our methylome-wide results.

### Colocalized *cis*-mQTL effects of non-MHC MS susceptibility loci

Of the 107,922 significant *cis*-mQTL effects, top mQTL SNPs for 3,090 CpGs were located within 100kb of the top non-MHC MS-associated SNPs^2^ (n=200). To determine the loci with likely shared causal effects on *cis* DNA methylation and MS susceptibility while minimizing false positive findings as a consequence of coincidental overlap because of linkage disequilibrium (LD), we performed Bayesian colocalization analyses^11^ between MS and *cis*-mQTL effects. MS susceptibility summary association statistics were taken from the discovery phase of the recent MS GWAS^2^ performed using data from 41,505 individuals. We found strong evidence of colocalization (posterior probability >0.8) between 19 MS-associated loci and 43 *cis*-mQTL effects (**Supplementary Table 3**). The top three colocalized loci (posterior probability >0.95) are shown in **Figure 2A**: the rs2248137 MS effect on chromosome 20 was colocalized with the *cis*-mQTL effect for cg14595058 upstream of the *CYP24A1* gene; the rs1077667 MS effect on chromosome 19 was colocalized with the *cis*-mQTL effect for cg23071186 located in the *TNFSF14* gene; and the rs7731626 MS effect on chromosome 5 was colocalized with *cis*-mQTL effects for 13 CpGs located in and around the *ANKRD55* and *IL6ST* genes (8 CpGs with posterior probability >0.95 and 5 CpGs with posterior probability >0.8). Further examination of the 13 CpGs in the chromosome 5 colocalized locus revealed that they were all located in methylation “open sea” regions spanning over an area up to 172kb from the top MS SNP. Reference chromatin state annotation (Roadmap Epigenomics Project^10^) suggested that they were located in putative enhancer regions, and their methylation levels were positively correlated with each other (**Supplementary Figure 2**). Colocalization with mQTL effects has previously been reported for this locus in the BLUEPRINT study^9^. The 16 other colocalized MS loci with posterior probability >0.8 and <0.95 affected methylation levels of 28 CpGs including CpGs in the vicinity of *C1orf106, RGS14, AHI1, CHST12, ZNF767P, TRIM14, SLC15A3, RMI2, TBX6, TEAD2, CCDC155, NCOA5* and *NCF4* genes (**Supplementary Table 3**).

**Figure 2.**
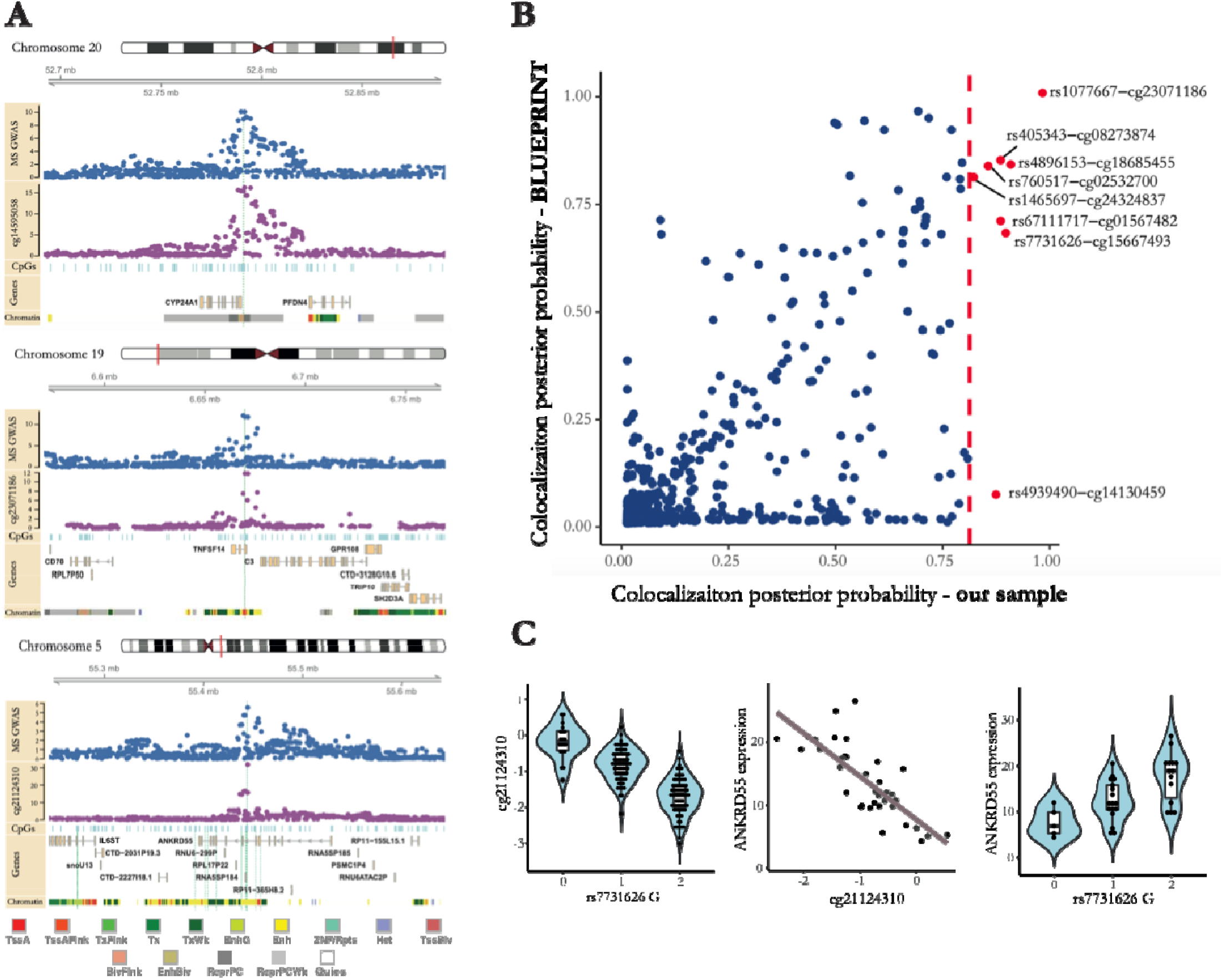
Colocalized *cis*-mQTL effects of MS susceptibility loci. **(A)** The top 3 colocalized MS-*cis*-mQTL effects are illustrated (posterior probability >0.95 for colocalization between MS susceptibility and ≥ 1 *cis*-mQTL effect). The MS GWAS rows (*blue* dots) show −log(*p*-value) of association between SNPs and MS susceptibility in the discovery phase of the 2019 IMSGC GWAS. CpG rows (*plum* dots) show −log(*p*-value) of association between SNPs and the specified *cis*-CpGs methylation levels. Locations of the local CpGs measured with the Infinium MethylationEPIC kit are shown using *light blue* vertical lines. Gene exon/intron positions are based on Ensembl 93. Chromatin state annotations for CD4^+^ T cells are downloaded from the Roadmap Epigenomics Project (sample #E043). *Green* vertical lines represent the genomic location of the colocalized CpGs: Long vertical lines traversing all rows represent the top specified CpGs, while other vertical lines represent the additional colocalized *cis* -mCpGs with posterior probability >0.8. All genomic positions are in GRCh37 (hg19) coordinates. **(B)** Comparison between MS-*cis*-mQTL colocalization posterior probabilities using mQTL summary statistics from our study and BLUEPRINT for the available common CpGs. Vertical *red* line represents the threshold for high colocalization posterior probability in our study (i.e. 0.8). **(C)** The *cis*-mQTL, *cis*-eQTL, and CpG-mRNA association for the chromosome 5 top MS-*cis*-mQTL colocalized effect. Methylation levels are shown in M-values. Gene expression values are in TPM.

**Figure 3.**
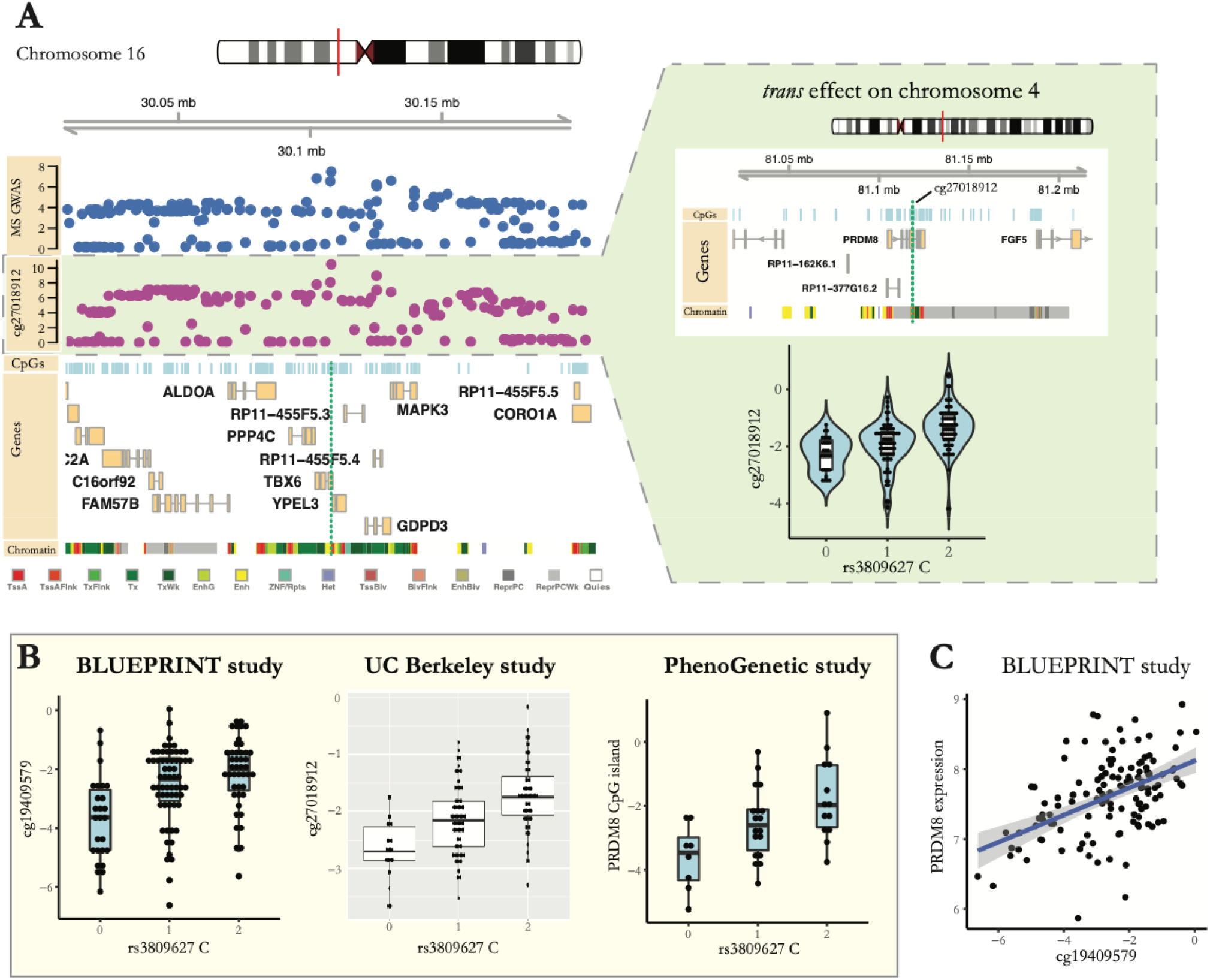
Colocalized *trans*-mQTL effect of MS susceptibility locus centered on rs3809627. **(A)** The top colocalized MS-*trans*-mQTL effect observed between MS susceptibility locus centered on rs3809627 in chromosome 16 and cg27018912 located in a CpG island in chromosome 4. *Left:* Association (-log(*p*-value)) between SNPs located in the chromosome 16 locus and MS susceptibility (*blue* dots) and the *trans*-mCpG cg27018912 methylation levels (*plum* dots), accompanied by visualization of the CpGs, genes and modeled chromatin states in the chromosome 16 locus. *Green* vertical lines represent additional colocalized *cis*-mCpGs (posterior probability >0.8). *Right:* Genomic position of the *trans*-mCpGs (cg27018912, cg19409579 and cg04235768) on chromosome 4 (vertical *green* lines) in relation to nearby CpGs, genes, and modeled chromatin states. The top colocalized *trans*-mQTL effect is shown in the *bottom* using methylation M-values. **(B)** Replication of the identified *trans*-mQTL effect between rs3809627 MS susceptibility variant (chromosome 16) and CpGs in the CpG island located on chromosome 4 in three independent studies. Methylation levels are shown in M-values. Data is shown for cg19409579 in the BLUEPRINT study as the top available *trans* -mCpG measured with the Infinium HumanMethylation450 array. Methylation data shown for the PhenoGenetic study are the average methylation levels of all the measured CpGs in the identified CpG island. **(C)** Significant CpG-mRNA association observed between cg19409579 *trans*-mCpG and *PRDM8* gene expression in the BLUEPRINT study. Methylation levels are in M-values. Gene expression levels are ComBat normalized values available from the BLUEPRINT.

We further compared the results of our colocalization analysis with the outcome of a similar analysis that we performed using *cis*-mQTL summary statistics obtained from the BLUEPRINT study. Data from 1,414 CpGs (out of 3,090) were shared between the two MS-*cis*-mQTL colocalization analyses, and high correlation was observed between colocalization posterior probability estimates (Spearman’s ρ= 0.71) (**Figure 2B**). Of the 43 CpGs with colocalized MS-*cis*-mQTL effects in our data, eight were available in BLUEPRINT. All, except for the cg14130459 *cis*-mQTL effect in the *SLC15A3* gene, showed replication, with comparable evidence of colocalization using the BLUEPRINT data (posterior probability estimates >0.67, **Supplementary Table 3, Figure 2B**). The lack of colocalization evidence between MS and the cg14130459 *cis*-mQTL effect might be explained by the weaker mQTL association observed for cg14130459 in naïve CD4^+^ T cells of healthy participants (*p*=0.006, BLUEPRINT study) in comparison to CD4^+^ T cells of MS patients (*p*=2.68×10^−10^, our study).

Taking advantage of available RNA sequencing data from primary naïve and memory CD4^+^ T cells from a subset of our subjects (n=36), we examined the gene expression correlates of the 43 identified CpGs with colocalized MS susceptibility and *cis*-mQTL effects. Our association analyses between methylation levels of the identified CpGs and expression levels of the genes in *cis* (±1Mb of each CpG) revealed significant inverse correlations between *ANKRD55* gene expression and *cis* CpG methylations at 5% FDR (top association: cg21124310: *p*= 2.7×10^−6^ in naïve and *p*= 1.2×10^−5^ in memory CD4^+^ T cells). The MS risk allele at this locus, rs7731626^G^, has previously been linked with higher *ANKRD55* expression, and the known *cis*-eQTL effect has been demonstrated to colocalize with the MS effect in CD4^+^ T cells of healthy individuals^5,9^. Similarly, we observed *cis*-eQTL effects for the MS SNP on *ANKRD55* expression in both naïve (*p*= 6.9×10^−4^, t= 3.7, **Figure 2C**) and memory CD4^+^ T cells (*p*= 1.0×10^−4^, t= 4.4) in our MS subjects. Our analyses further suggested that the effect of rs7731626^G^ on *ANKRD55* expression is mediated by its effect on cg21124310 methylation (*p* for mediated effect <2×10^−16^ and *p* for direct effect >0.05 in both naïve and memory CD4^+^ T cells; also *p* for mediated effect of *ANKRD55* expression on cg21124310 methylation >0.05). We note that our sample size for the DNA methylation-gene expression sub-analysis was small, limiting our power to identify the weaker gene expression correlates of other colocalized MS-mQTL effects. However, the identified genetic loci and their corresponding CpGs can be used in future studies of proper sample size with specific focus on identifying the gene expression correlates of these loci and their relationship with methylation effects.

We also reviewed the *cis*-eQTL studies of peripheral blood mononuclear cells^2^, purified blood immune cells^3,6,9^ and brain tissue^8^ from MS patients or healthy controls, and summarized the results for top MS and top mQTL SNPs of the colocalized MS-*cis*-mQTL effects in **Supplementary Table 4**. We note that a number of our colocalized loci - namely MS SNPs in the vicinity of *C1orf106, CHST12, CXCR5, CYP24A1, LINC01967, NCOA5, SOX8* and *TNFSF14* genes - have not previously been associated with *cis*-eQTL effects in CD4^+^ T cells and need to be investigated further for their functional effects.

### Colocalized *trans*-mQTL effects of non-MHC MS susceptibility loci

To investigate the influence of MS susceptibility loci on DNA methylation at distant CpG sites (*trans*-mQTL effects), we first mapped the *trans*-mQTL effects of all variants within 100kb of the top MS-associated SNPs. We then performed colocalization analysis and found evidence for colocalization (posterior probability >0.95) of the effect of the MS locus centered on rs3809627 (chromosome 16) and 3 *trans*-mQTL effects. All three *trans*-mCpGs were located in a CpG island on chromosome 4 in the area flanking a transcription start site in the *PRDM8* gene (*p*-value for the top colocalized mQTL effect: rs3809627-cg27018912= 5.7×10^−11^, t= 7.1, colocalization posterior probability= 0.98) (**Figure 3A, Supplementary Table 3**). While this MS locus centered on rs3809627 also showed colocalization with 2 *cis*-mQTL effects (CpGs in an enhancer region in the *TBX6* gene; colocalization posterior probability= 0.92), the methylation levels of the *trans* and *cis* mCpGs were not correlated (**Supplementary Figure 3**), suggesting that the *trans*-mQTL effects of rs3809627 were not directly mediated through its *cis*-mQTL effects.

We replicated our *trans*-mQTL finding - rs3809627 on chromosome 16 and the chromosome 4 CpG island - in three independent datasets (**Figure 3B**). First, we found the same effect using the BLUEPRINT naïve CD4^+^ T cell DNA methylation data (*p*=4.8×10^−6^). Second, we replicated the *trans*-mQTL in a dataset of DNA methylation profiles from whole blood of 208 MS patients from UC Berkeley (*p*=5.4×10^−13^). Third, we analyzed targeted methylation data from CD4^+^ T cells purified from 48 healthy participants in the PhenoGenetic study^12^ (*p*=7.1×10^−4^) (see **Methods** for more details). These results generate high confidence in the *trans*-mQTL effect of rs3809627 and establish it as an effect seen in both MS and healthy subjects. Given the previous implications of activated CD4^+^ T cells in the pathophysiology of MS, we had generated, in parallel, a dataset of *ex vivo* activated CD4^+^ T cells using a subset of the PhenoGenetic subjects (n=28) to investigate the effect of *ex vivo* activation on the methylation of CpGs of interest. We did not observe a significant difference in the CpG island methylation in the pairs of non-stimulated and stimulated samples (*p*=0.95).

Assessing the possible role of these *trans*-mCpGs, we observed a highly significant positive association between the target *trans*-mCpG (cg19409579) methylation level and *PRDM8* mRNA expression in data from naïve CD4^+^ T cells in the BLUEPRINT study (n=127, ρ=0.49, association *p*-value adjusted for age and sex= 4.7×10^−9^, **Figure 3C**). PRDM8 belongs to a family of histone methyltransferases that are believed to act as negative regulators of transcription and play an important role in development and cell differentiation. Hypermethylation of *PRDM8* is associated with the first step of thymocyte differentiation (transition from thymic progenitors with lymphomyeloid potential to T-lineage-restricted progenitors)^13^. Among T cell subtypes, *PRDM8* has higher expression levels in regulatory, memory and *ex vivo* activated CD4^+^ T cells^6^. Its chromatin accessibility increases upon stimulation in naïve CD4^+^ T cells, and the region remains accessible in memory CD4^+^ T cells^14^. Although *PRDM8* has not previously been associated with MS, our replicated *trans*-mQTL findings suggests a role for this gene in the peripheral immune pathophysiology of MS as a downstream effect of the rs3809627 MS susceptibility variant.

### *cis* and *trans* DNA methylation effects of an MS susceptibility polygenic score for the MHC

The MHC has a complex long-range LD structure and harbors 32 independent genome-wide significant associations with MS^2^, including the haplotype with the largest MS-associated risk, *HLA-DRB1*1501*. Given the region’s complexity, we elected to focus our investigation on an assessment of the summary risk of MS based on the susceptibility variants found within the MHC. Specifically, we calculated a weighted MHC polygenic score representing the aggregate genetic effects of all of the independent loci in this region and used this measure as the outcome for our methylome-wide association study (MWAS). Our MWAS revealed significant associations between the MS MHC polygenic score and the methylation of 55 CpGs (FDR <0.05), 54 of which were located inside the extended MHC region on chromosome 6 and one was located on chromosome 17 (**Figure 4A, Supplementary Table 5, Supplementary Figure 4**).

**Figure 4.**
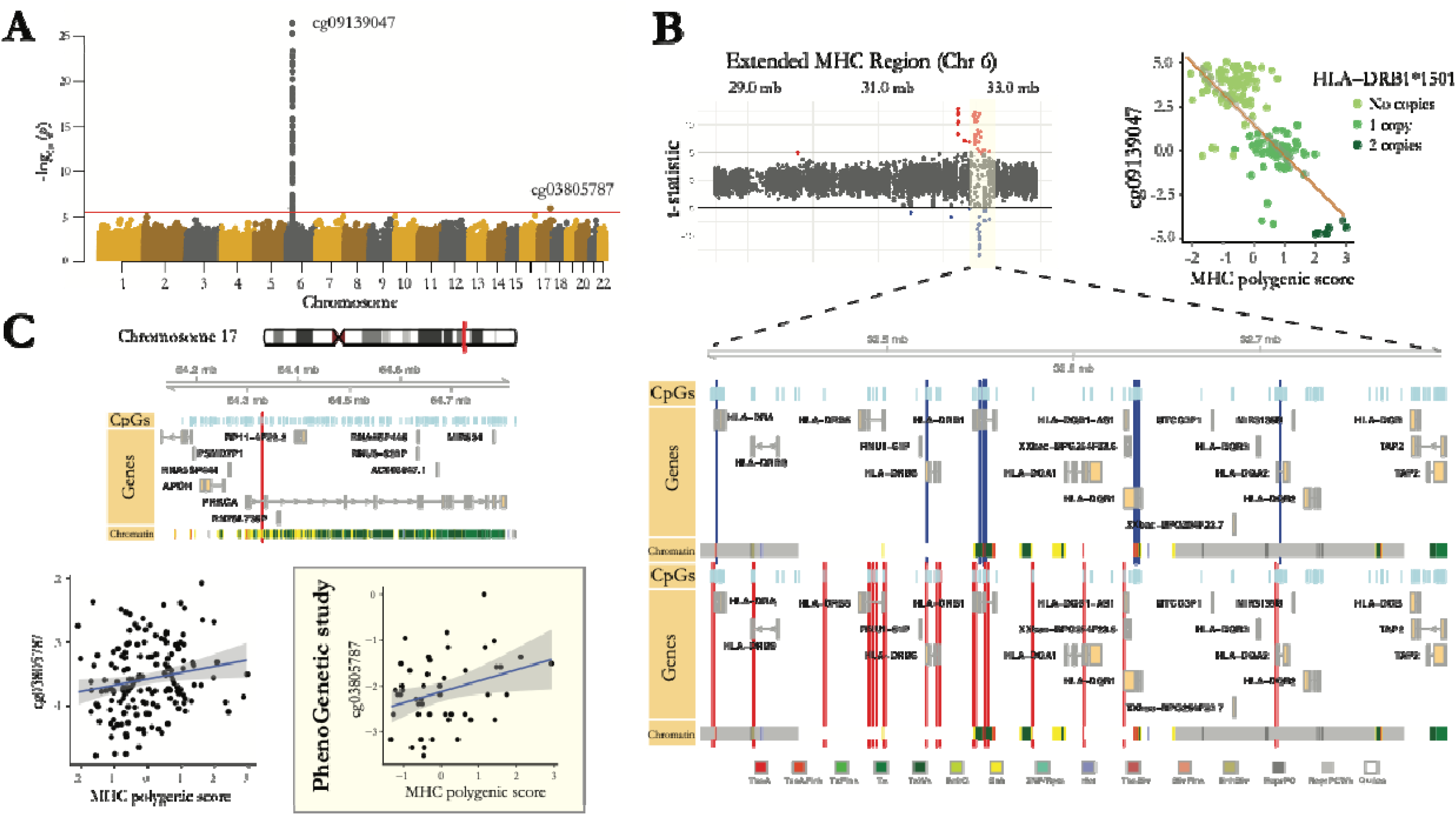
*cis* and *trans* DNA methylation effects of MS MHC polygenic score. **(A)** Manhattan plot shows association *p*-values between CpG methylation levels and MS-associated MHC polygenic score. *Red* horizontal line shows the significance threshold (FDR<5%). Top CpGs in the two significant regions are specified. **(B)** *t*-statistics for positive and negative associations between the methylation levels of the CpGs located inside the extended MHC region and the MHC polygenic score are shown (*red:* positive, *blue:* negative). The inset illustrates the genomic location of the significant CpGs in the MHC class II region in relation to nearby CpGs, genes, and modeled chromatin states. *Blue*/*red* vertical lines represent the locations for CpGs with negative/positive associations with the MHC polygenic score, respectively. Association between the top identified CpG, cg09139047 located in the *HLA-DRB1* gene, and MHC polygenic score (shown in *Z*-scores) is visualized in the *top right* corner. Colors represent the number of *HLA-DRB1*1501* alleles. **(C)** The genomic location for cg03805787 (associated with the MHC polygenic score in chromosome 17, *red* vertical line) is shown in relation to surrounding CpGs, genes, and modeled chromatin states. *Bottom* panel illustrates the association between cg03805787 methylation levels (M-values) and MHC polygenic score (*Z*-scores) in our main study of MS subjects, as well as the replication in the healthy subjects of the PhenoGenetic study.

The 54 identified CpGs located in the MHC region consisted of both hypo- and hyper-methylated CpGs in association with a higher MS MHC polygenic score (**Figure 4B, Supplementary Figure 5**). The majority of these CpGs were located in the MHC class II region, which also included the most significant association (a CpG located in the *HLA-DRB1* gene). The majority of the hypo-methylated CpGs were located in two CpG islands in or around the transcription start sites of *HLA-DRB1* and *HLA-DQB1* genes, although both genes also had a few hyper-methylated CpGs. On the other hand, the majority of the hyper-methylated CpGs were located in *HLA-DRB5, HLA-DRB6* and *LOC101929163 (XXbac-BPG154L12.4*). Our findings are supported by previous studies^15,16^ which also observed hypo- and hyper-methylated CpGs in the MHC class II region in MS patients in comparison to healthy controls and also in association with the *HLA-DRB1*1501* genotype. Moreover, replication analysis using the BLUEPRINT data showed compelling evidence of association with the MHC polygenic score for all available CpGs (**Supplementary Table 5**).

The only CpG significantly associated with the MS MHC polygenic score outside of the MHC region was located in an enhancer of the *PRKCA* gene on chromosome 17 (cg03805787, *p*= 1.2×10^−6^, *q*= 0.018, t= 5.8) (**Figure 4C**).The effect of the MHC polygenic score on cg03805787 methylation was not mediated by the methylation of the CpGs in the MHC region. Additionally, this effect was not explained only by the *HLA-DRB1*1501* genotype, as the effect of the MHC polygenic score remained significant (*p*= 0.002) after accounting for the effect of *HLA-DRB1*1501* in the model. Replication analysis for cg03805787 could not be performed using data from the BLUEPRINT study, as measurement of cg03805787 methylation is not included in the Illumina 450K array. However, we did replicate the association of the MHC polygenic score and cg03805787 methylation in CD4^+^ T cells from healthy participants in the PhenoGenetic study (*p*=0.025, **Figure 4C**). No significant difference was found in cg03805787 methylation levels between primary and *ex vivo*-activated CD4^+^ T cells in these individuals (*p*=0.84). This finding on the effects of MHC genetic variation on DNA methylation in the *PRKCA* region opens an interesting arena for further investigation, as this gene may play a role as a focal point for the functional consequences of MHC variants.

### DNA methylation effects of MS susceptibility total polygenic score

In our final analysis, we investigated the effects of the aggregate genetic risk score for MS (i.e. the sum of the MS MHC and non-MHC polygenic scores) on DNA methylation in CD4^+^ T cells through an MWAS study. 38 CpGs showed significant association with this MS total polygenic score at FDR<0.05 (**Figure 5A, Supplementary Table 6**). 36 of these CpGs were located in the MHC and were also identified in association with the MS MHC polygenic score alone. The non-MHC polygenic score did not contribute to these associations. In contrast, the association of 2 CpGs that were found outside of the MHC, on chromosomes 10 and 17, could not be explained by *cis* effects. Methylation of cg16050799 (in an intron of the *CRHR1* gene on chromosome 17, **Figure 5A**) was affected by independent and additive effects of both MHC (*p*= 7.6×10^−4^) and non-MHC (*p*= 1.9×10^−6^) polygenic scores, suggesting a convergence of the effects of multiple MS-associated loci on this methylation site. On the other hand, cg19223119 methylation (located in the *DIP2C* gene on chromosome 10) was mainly associated with the MHC polygenic score (*p* for MHC score= 6.5×10^−6^, *p* for non-MHC score= 0.024).

**Figure 5.**
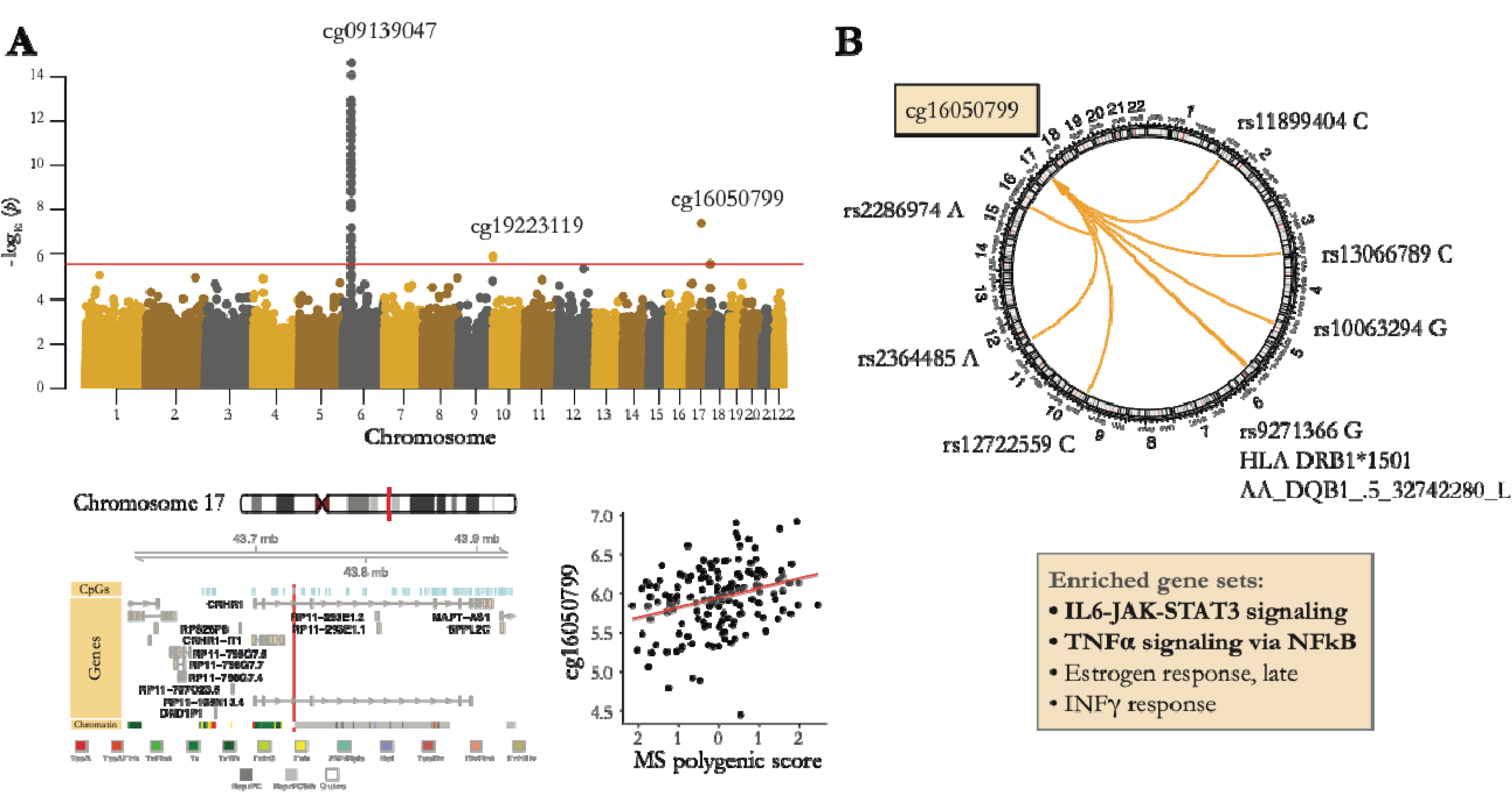
DNA methylation effects of MS total polygenic score. **(A)** Manhattan plot shows association *p*-values between CpG methylation levels and MS total polygenic score. *Red* horizontal line shows the significance threshold (FDR<5%). Top CpGs in the three significant regions are specified. The genomic location for cg16050799 (the top associated CpG outside the MHC region, *red* vertical line) in relation to nearby CpGs, genes, and modeled chromatin states, in addition to its association with the MS total polygenic score (*Z*-scores) is further shown. Methylation levels are shown in M-values. **(B)** Genomic locations of MS susceptibility alleles nominally associated with cg16050799 methylation (*p*<0.05) are shown. Gene sets enriched for genes associated with these loci are specified.

As cg16050799 methylation was affected by both MHC and non-MHC polygenic scores, we decided to further characterize this effect and identify the individual MS-associated loci which were driving this association. Although no individual association passed the multiple comparison testing correction (FDR <0.05), results revealed suggestive (*p* <0.05) associations between 3 MHC and 6 non-MHC risk loci and higher cg16050799 methylation. Gene set enrichment analysis performed on genes associated with the 6 non-MHC risk loci showed enrichment for the IL-6 and NF-kB pathways (**Figure 5B, Supplementary Table 7**). Replication analysis could not be performed for cg16050799, as the methylation measurement was not available as part of the BLUEPRINT methylation array and the measurements performed in the PhenoGenetic samples did not pass quality control. In addition, the MHC variants were not imputed in the UC Berkeley study, and the association between non-MHC polygenic score and cg16050799 methylation was not significant in their whole blood sample. Given the shortcomings of our replication attempts, we could not determine the replicability of the observed association. However, we believe that convergent effect of both MHC and non-MHC polygenic scores on cg16050799 methylation in CD4^+^ T cells of MS patients deserves more thorough investigation in future studies. CRHR1 is a receptor protein for corticotropin releasing hormone (CRH). Peripheral CRH is believed to have proinflammatory effects, with CRH-deficient mice showing an attenuated form of EAE and their CD4^+^ T cells showing reduced proliferation and a shift towards Th2 cytokine profile^17^. Our finding, if replicated, might provide support for the role of corticotropin hormone-related pathways in MS treatment or prevention.

## DISCUSSION

Our examination of the genetic architecture of DNA methylation profiles of primary CD4^+^ T cells from MS patients yielded two major products. First, we provide a new **Resource**: a genome-wide *cis*-mQTL map for primary CD4^+^ T cells which is more extensive than the existing map (e.g. the *cis*-mQTLs available from the BLUEPRINT Epigenome Project^9^) due to a greater number of DNA methylation sites being assessed. Our extended T cell DNA methylation mQTL map will therefore be useful to a wide variety of investigators, including those outside of the field of MS. Results for the significant *cis*-mCpGs are available in **Supplementary Table 2**. Importantly, there is high concordance of the *cis*-mQTL effects in CD4^+^ T cells from MS patients (our study) with those found in naïve CD4^+^ T cells from healthy individuals (the BLUEPRINT study)^9^ (**Figure 1B**), which (1) increases our confidence in the robustness of both resources and (2) suggests that the majority of our *cis*-mQTL effects are not specific to MS. This makes our *cis*-mQTL map an excellent exploratory resource for the study of *cis* functional effects of genetic loci associated with other diseases that have CD4^+^ T cell involvement and other CD4^+^ T cell-related traits such as CD4^+^ T cell functional phenotypes.

Second, in a series of analyses focused on MS, we identified not only the *cis* effects of individual MS susceptibility loci but also robust evidence for *trans* effects: (a) the MS locus near *TBX6* on chromosome 16 influences methylation of a CpG island in the *PRDM8* gene on chromosome 4, and (b) the MHC polygenic score affects methylation of an enhancer of *PRKCA* on chromosome 17. Both of these results are supported by replication analyses performed using independent datasets of healthy subjects and MS patients. Such robust *trans-QTL* effects are notoriously difficult to identify, and, here, they suggest that there may be convergence of the effects of multiple MS risk variants on DNA methylation of sites that are not themselves genetically implicated in MS. Given that the MHC is by far the largest genetic risk factor for MS, the *PRKCA* region may offer a potential alternative to functional dissection of the genetically complex MHC region as we develop prevention strategies. The *PRKCA* locus has been linked genetically with MS in familial MS cases in independent sets of British^18^, Finnish and Canadian families^19^. Protein kinase C alpha (PKCα, encoded by the *PRKCA* gene) is a member of a family of serine/threonine protein kinases which is expressed ubiquitously in T cells and is involved in T cell receptor comodulation (i.e. downregulation of non-engaged T cell receptors following peptide-MHC complex engagement and T cell activation)^20^. Programmed downregulation of T cell receptors is believed to represent a negative feedback mechanism that constrains T cell effector function in order to avoid excess inflammatory damage^21^. In addition, PKCα has been implicated in signaling pathways necessary for T cell interferon gamma^22^, interleukin (IL)-2^23^ and IL-17A^24^ production which are implicated in MS pathogenesis, and Prkca deficiency protects mice from experimental autoimmune encephalomyelitis (EAE, an MS animal model)^24^. Finally, pharmacologic inhibition of PKCα has been shown to be beneficial in models of a number of other T cell-related immune conditions^25^. This extensive biological understanding of PRKCA is consistent with our observation that the effects of the MHC polygenic score on MS susceptibility may be exerted, in part, through this locus. Likewise, our finding on the effect of MS total polygenic score on methylation of the *CRHR1* region, if replicated, would provide an interesting target for modulation of the convergent effects of several MS susceptibility loci.

In terms of the 19 MS susceptibility loci with colocalized *cis*-mQTL effects (with the *TBX6* locus also having the colocalized *trans*-mQTL effect), seven were confirmed in the BLUEPRINT data, and the other 12, while significant in our analysis, will need further validation. While these analyses prioritize certain haplotypes as likely having effects on both MS susceptibility and *cis* or *trans* DNA methylation, colocalization methods are limited in their ability to distinguish between two colocalized effects and two independent effects whose causal variants are in very high LD. Moreover, colocalization does not necessarily represent a mediatory causal relationship whereby one of the traits (e.g. DNA methylation) mediates the effect of genetic variation on the other trait (e.g. MS risk). A pleiotropic effect (i.e. independent effects of genetic variation on DNA methylation and MS risk) is alternatively possible, and the exact nature of the relationship (mediation, partial mediation, or full independence) cannot be resolved without large-scale longitudinal data. Nevertheless, colocalized MS and mQTL effects are reasonable candidates for further molecular and functional interrogation.

In addition, we acknowledge that our DNA methylation measurements were performed only on primary CD4^+^ T cells and not in the other cell types known to be involved in MS pathophysiology (such as B cells, NK cells, monocytes and microglia). Moreover, due to the cross-sectional design of our study, we were not able to take into account the dynamics of DNA methylation. Finally, our analyses are restricted to those CpGs interrogated by the array.

Although available information from previous eQTL and protein QTL studies have helped to identify the functional consequences of a number of MS risk variants^2–5,26^, it is notable that not all of our identified DNA methylation-associated MS variants have previously been associated with gene or isoform expression changes (**Supplementary Table 4**). This lack of gene level information can stem from a number of reasons, including cell type- or context-specificity of the effects, or exclusion of genes with lower levels of total mRNA or isoform expression from prior analyses. In addition to providing complementary information to eQTL results, mQTL studies can be used to prioritize genetic loci for more thorough investigation of their gene level effects. In line with this, and despite the moderate sample size, we were able to show that the rs7731626 MS variant’s effect on *ANKRD55* RNA expression is likely caused by altered DNA methylation in *cis*.

Our new data **Resource** has therefore generated clear paths for further investigating the molecular events that contribute to the onset of MS at 19 of the more than 150 validated susceptibility loci. By sampling CD4^+^ T cells in an inflammatory disease population, we have not only validated the robustness of the BLUEPRINT results for this cell population but also made the important observation that most of the evaluated mQTLs are not significantly different in health and disease. This makes our expanded set of 107,922 (82,374 new) CD4^+^ T cell mQTLs of interest to a broad array of investigators interested in this important cell type outside of the MS context. Finally, our strategy of targeted *trans*-mQTL investigations returned replicable results in the *PRDM8* and *PRKCA* loci, providing important evidence of the propagation of the functional consequences of MS variants outside of their immediate vicinity and a demonstration of the utility of our approach to find distal functional consequences. Our results will therefore help to ground future study designs to continue to elaborate the cascade of events leading to MS onset.

## METHODS

### Main methylation study

#### Subjects

Subjects were participants in the Comprehensive Longitudinal Investigation of Multiple Sclerosis at the Brigham and Women’s Hospital (CLIMB) study^7^. CLIMB is a natural history observational study of MS, in which participants undergo semi-annual neurological examinations and annual magnetic resonance imaging and blood draw, from which peripheral blood mononuclear cells (PBMC) are cryopreserved. In 2015, cryopreserved PBMC samples from subjects meeting the following criteria were pulled from the archive for sample processing: (1) age 18-55 years old, (2) a diagnosis of MS fulfilling 2010 McDonald criteria, (3) a relapsing-remitting disease course at the time of sampling, (4) being on disease-modifying therapies (either glatiramer acetate [GA] or dimethyl fumarate [DMF]) at the time of sampling, (5) no evidence of disease activity in the prior 6 months, (6) no steroid use in the preceding 30 days, and (7) an Expanded Disability Status Scale (EDSS) score between 0 to 4 at the time of sampling. The study was approved by the Institutional Review Board of the Brigham and Women’s Hospital, and all participants had signed a written informed consent to participate in CLIMB. Samples from 208 patients who fulfilled the criteria were used for further processing.

#### Peripheral blood CD4^+^ T cell isolation

Peripheral blood mononuclear cells (PBMC) were collected prospectively from participants using the Immune Tolerance Network (https://www.immunetolerance.org/) protocol to ensure data quality and to minimize variation among samples. In short, fresh blood was processed within 4 hours using the Ficoll procedure to extract PBMC. The PBMC were then resuspended in fetal bovine serum with 10% DMSO and cryopreserved in liquid nitrogen. The samples meeting study criteria were removed from the sample archive and thawed in batches. Cell viability was accessed by acridine orange/propidium iodine staining and CD4^+^ T cells were isolated using a positive selection strategy implemented with Miltenyi magnetic CD4 MicroBeads (130-045-101). 1.7 million CD4^+^ T cells were obtained from each vial (range: 0.26-6.9 million). Isolated cells were resuspended into lysis buffer, and DNA was extracted from each sample using QIAamp DNA Blood Mini Kits (Qiagen 51104).

#### DNA methylation data

Whole-genome DNA methylation measurement was performed on DNA from purified CD4^+^ T cells using Infinium MethylationEPIC BeadChip (Illumina 850K array) by the Center for Applied Genomics Genotyping Laboratory at the Children’s Hospital of Pennsylvania. Quality control was performed by comparing log2 median intensities of methylated and unmethylated channels, inspecting samples’ beta distributions and control probes bisulfite conversion rates, and removing samples with detection *p*-values >0.05 at >1% of the methylation sites. No sample was excluded at this step and average probe detection *p*-values were <0.01 for all samples. Further processing included normalization using Noob^27^ and BMIQ^28^ algorithms, removing methylation sites with beadcount <3 in >5% of samples or detection *p*-value >0.01 in any sample, in addition to sites located on sex chromosomes, sites associated with probes with polymorphic targets with minor allele frequency >1% in individuals with European ancestry^29^, and sites associated with cross-reactive probes^29^. M-values were calculated for 769,699 methylation sites to be used in statistical analysis. Data from probes measuring single nucleotide polymorphisms were also extracted to be used for identity check by cross-referencing with the genetic data.

#### Genotyping data

Whole-genome genotyping data were available for the majority of participants (n=180) as part of previously genotyped batches of CLIMB samples using Illumina MEGA-EX and Affymetrix 6.0 arrays. SNPs with call rate <95% and samples with genotyping rate <90%, mismatch between recorded sex and genetic sex, and low or high heterozygosity rates (>3 standard deviation) were removed. Data from the two genotyping batches were merged and samples from related individuals (pi-hat >0.125) were also excluded. Multidimensional scaling was performed using HapMap3 reference data. Samples from individuals of non-European ancestry and ethnic outliers with >3 standard deviation difference from the European samples were identified and further removed (final N at this stage= 158). Imputation was performed for each genotyping batch separately using the Michigan Imputation Server^30^ and the Haplotype Reference Consortium (HRC) panel v1.1. Imputed data from the study participants were merged, and SNPs with low imputation quality (R^2^ <0.8) were removed.

#### Final dataset

Principal component (PC) analysis was performed on the methylation data from individuals that passed genetic data quality control (n=158). The first four principal components explained 39% of the total variability (**Supplementary Figure 6**) and were used to remove outlier samples from the methylation data (>3 standard deviation on any of the 4 PC values). Two additional samples were removed using this measure, and the final dataset consisted of data from 156 individuals (demographic details are available in **Supplementary Table 1**).

#### mQTL analyses (*cis* and *trans*)

We used QTLtools^31^ to perform genome-wide mapping of CD4^+^ T cell *cis*-mQTLs, as well as targeted mapping of *trans*-mQTLs among SNPs in 100kb of the top autosomal non-MHC MS-associated SNPs reported in ^2^. SNPs with minor allele frequency <0.05, minor allele count <15, and departure from Hardy-Weinberg equilibrium (*p* <1×10^−6^) were excluded from mQTL analyses.

For the *cis*-mQTL analysis, linear regression analysis was performed between quantile normalized DNA methylation M-values and imputed genotype dosages for SNPs in *cis* (±1Mb) of each CpG site, accounting for the effects of age, sex, treatment, genotyping array, the first 3 genotyping PCs and the first 4 methylation PCs. The nominal *p*-values for the top variants in *cis* were then adjusted for the number of tests performed in *cis* using a permutation scheme (n=1000 permutations) which models the null distribution of associations using a beta distribution. Finally, to account for the multiple CpG sites tested across the whole genome, significant associations were reported at 5% False Discovery Rate (FDR).

For the *trans*-mQTL analysis, linear regression analysis was performed between SNPs in ±100kb of the top MS-associated SNPs and all CpG sites outside their *cis* windows, accounting for the effects of the previously mentioned covariates. The nominal *p*-values were adjusted for the number of variants being tested using the null distribution of associations built from beta approximation of permutation outcome by permuting and testing associations between randomly selected 10,000 CpG methylation levels and all included variants. FDR adjustment was then implemented to account for the multiple CpG sites being tested.

#### Colocalization analyses

Analyses were performed using the R “coloc” package^11^ in order to determine whether mQTL effects and MS susceptibility effects that are located in close proximity of each other are likely the result of a shared causal variant. Briefly, for each autosomal non-MHC MS locus, mQTL effects whose top SNPs were located in 100kb of the top MS SNP were identified. Colocalization of the effects of each identified MS-mQTL pair was assessed separately. The input data for each analysis consisted of: 1) summary association statistics (*p*-value, beta -regression coefficient-, variance of beta, sample size, and minor allele frequency) of SNPs in 100kb of the top MS SNP extracted from the discovery phase of the IMSGC GWAS study^2^ (n=14,802 MS and 26,703 control participants), as well as our *cis* or *trans* mQTL analyses; 2) the suggested prior probabilities^11^ of 1×10^−4^ for the association between each SNP and each of the two traits, and 1×10^−6^ for the association between each SNP and both traits. The posterior probability of each locus containing a causal variant affecting both MS and mQTL effects was estimated against the probabilities of other models (a null model of no association, association with only the first or the second trait, or independent associations with each of the two traits). Results are reported at a posterior probability cut off of 0.8 for the colocalized effects.

#### Polygenic score calculations

MS polygenic scores were calculated based on the genome-wide significant results from the 2019 IMSGC MS GWAS study (31 MHC markers and 200 autosomal non-MHC SNPs)^2^. SNPs, amino acids and HLA alleles in the MHC region were imputed using the “SNP2HLA” package^32^ and the Type 1 Diabetes Genetics Consortium (T1DGC) HLA reference panel. The MS MHC polygenic scores were then calculated as the sum of the imputed dosages of the 31 MHC markers associated with MS multiplied by their effect sizes^2^ (log odds ratio). For the MS total polygenic score, SNP dosages for the autosomal non-MHC MS SNPs (or if not available, tag SNPs in LD >0.8) were extracted from the wholegenome imputed data (imputed using the Michigan Imputation Server^30^ and the HRC reference panel), multiplied by their effect sizes, and added to the MHC polygenic score. Data from 10 of the 200 non-MHC SNPs were not available in the quality controlled imputed data and were substituted by a close by SNP in LD >0.8. Data from 7 SNPs were not included in the MS total polygenic score, as no data were available for them or SNPs in LD >0.8 with them. The list of MHC and non-MHC markers used in the calculation of the MS total polygenic score and their effect sizes can be found in **Supplementary Table 8**.

#### Methylome-wide association studies

GLINT^33^ was used to perform the methylome-wide association studies (MWAS). The final methylation dataset was used to assess the associations between DNA methylation and MS polygenic scores at each of the methylation sites, while adjusting for the effects of age, sex, treatment, genotyping batch, the first 3 genotyping PCs and the first 4 methylation PCs. Significant results were reported at 5% FDR.

#### Gene set enrichment analysis

We used FUMA^34^ (v1.3.3d) for the gene set enrichment analysis. Briefly, Genotypes were mapped to genes (Ensembl Release 92) by positional (100kb window) and eQTL mapping using the “SNP2GENE” function, and the mapped genes were then fed to the “GENE2FUNC” function for the pathway analysis. Minimum overlap with each gene set was set to 2 genes. Results were reported at 5% FDR.

#### Statistical analysis

Post-hoc linear regression studies were performed while accounting for the effects of age, sex, treatment, genotyping batch, the first 3 genotyping PCs and the first 4 methylation PCs, unless otherwise specified. Enrichment analyses for chromatin states were performed using Chi-square tests. Mediation analyses were performed using R “mediation” package. All reported *p*-values are two-sided. All results are reported in GRCh37/hg19 genomic coordinates.

#### RNA sequencing sub-study

RNA sequencing data from peripheral blood purified naïve and memory CD4^+^ T cells were available for a subset of participants as part of a separate RNA sequencing study (n=42 and 43 naïve and memory CD4^+^ T cell samples, respectively). Data from participants who had changed medication between the sampling dates for DNA methylation and RNA sequencing studies were excluded (n=6 from each cell type). All remaining RNA sequencing samples were from patients receiving glatiramer acetate at the time of sampling for both studies. All RNA sequencing samples were taken within 5 years of the sampling date for the DNA methylation study, with 80% being performed within one year of the DNA methylation sampling. PBMC sampling and CD4^+^ T cells isolation were performed using the same protocol used in the DNA methylation study. Naïve and memory CD4^+^ T cells were isolated on a BD FACSAria flow cytometer using fluorochrome labelled CD3, CD4, and CD45RA antibodies. Whole transcriptome 25-bp paired end sequencing was performed using the Broad Institute HiSeq 2500 platform to an average depth of 15 million reads. Processing was performed according to the Broad Institute RNA-seq pipeline for the GTEx Consortium. Briefly, RNA sequence reads were aligned to the GRCh38/hg38 genome reference using STAR^35^, quality control was performed using RNA-SeQC^36^ and quantification of gene expression levels were performed using RSEM^37^. Transcripts with low expression values (average TPM <2) were removed. Samples with outlier average correlation with the other samples (|*D* statistic^38^| > 3 standard deviation from the mean) were excluded from further analysis (one memory CD4^+^ T cell sample was removed). TPM values were log-transformed and quantile-normalized. Gene start and end positions were extracted from Ensembl Release 93 annotations. Linear regression modeling was used to investigate the associations between methylation levels of CpGs of interest -identified from *cis*, *trans*, and polygenic score analyses- and mRNA expression levels of genes within ±1 Mb of their respected CpGs, while adjusting for the effects of age and sex. In total, 912 association analyses were performed for naïve CD4^+^ T cells assessing associations between 49 CpGs and 488 genes, and 887 association analyses were performed for memory CD4^+^ T cells between 49 CpGs and 480 genes, each using data from 36 participants. Associations with FDR-adjusted *p* <0.05 were considered significant.

## Additional datasets

### BLUEPRINT study

BLUEPRINT Epigenome Project’s publicly available *cis*-mQTL summary statistics from the study of naïve CD4^+^ T cells were used for comparison with our *cis*-mQTL and colocalization findings. DNA methylation measurements were performed using Illumina Infinium HumanMethylation450 BeadChips, and samples were from 132 healthy individuals of European origin (**Supplementary Table 1**). In order to perform replication analysis for our *trans*-mQTL and polygenic score analyses findings, we downloaded the BLUEPRINT genetic data from the European Genome-phenome Archive and performed whole-genome and MHC imputation of the data in a similar manner to the main methylation study. Associations were performed between genetic measures and BLUEPRINT’s publicly available processed DNA methylation data (M-values), accounting for the effects of age and sex.

### UC Berkeley study

Data from this study were both gathered and analyzed independently by the UC Berkeley investigators. Whole blood samples from 208 self-identified white, female MS patients were used for the study (**Supplementary Table 1**). Genome-wide DNA methylation was profiled using Infinium MethylationEPIC BeadChips. Methylation data were analyzed using Bioconductor “minfi” package^39^. Background dye correction and quantile normalization were performed, followed by the removal of batch effects using the ComBat^40^ function of R “sva” package. Ancestry and cell type heterogeneity were estimated using GLINT^33^, and methylation M values were adjusted for ancestry and cell type components as well as for age at sampling. Genome-wide genotyping was performed using Illumina Infinium 660K OmniExpress or OmniExpressExome BeadChip arrays. Merged 660K and Omni Express genotyping dataset (273,906 SNPs) were phased using SHAPEIT2, and imputed against reference haplotypes from Phase 3 of the 1000 Genomes Project using IMPUTE4. Linear regression models were used to study the association between adjusted M values and genotypes, while covarying for MS treatment status (0/1, only 13 patients were not receiving treatment). MHC imputation was not available for this study.

### PhenoGenetic study

Frozen PBMC samples from 48 healthy participants of European origin from the PhenoGenetic Project who were part of the ImmVar study^3,12^ and had available genotyping data were selected for replication analyses (**Supplementary Table 1**). CD4^+^ T cells were purified from cryopreserved PBMC using a magnetic microbead-based strategy (Miltenyi 130-096-533) after thawing with 10 mL PBS. Cells from a subset of the samples (n=28) were split in half, and one half was placed in culture with serum-free X-Vivo medium and then stimulated using 2 μg/mL anti-CD3 and 1 μg/mL anti-CD28 antibodies. Cells were collected 24 hours after stimulation. DNA was extracted from the 48 primary and 28 *in vitro* activated CD4^+^ T cell samples and underwent bisulfite conversion. Methylation levels of CpGs of interest were measured using the Agena Bioscience mass spectrometry-based EpiTYPER assay, with primers designed using EpiDesigner (**Supplementary Table 9**). Genotyping was performed as part of two larger batches of samples previously genotyped using Illumina Infinium OmniExpressExome and Illumina MEGA-EX arrays. Whole-genome and MHC imputation were performed as mentioned in the main methylation study. Association analyses were performed between genetic measures and DNA methylation M-values, adjusting for the effects of age and sex. Paired *t*-tests were used to compare the methylation levels between primary and activated cells at each CpG site.

## Data sharing

The complete set of summary statistics for the *cis*-mQTL analysis are searchable and available for download after the requester submits a request through our web form: https://docs.google.com/forms/d/e/1FAIpQLSdywPgMC3281y5aZJP8KRXhZqN7bXa9OQ9kaDpErBQ9C0-lTg/viewform

## Acknowledgement

We thank the participants of the CLIMB and PhenoGenetic studies for their contribution of blood samples from which the data were derived. The study was funded by a grant to PLD from the Massachusetts Life Sciences Center.

---

**Supplementary Figure 1.**
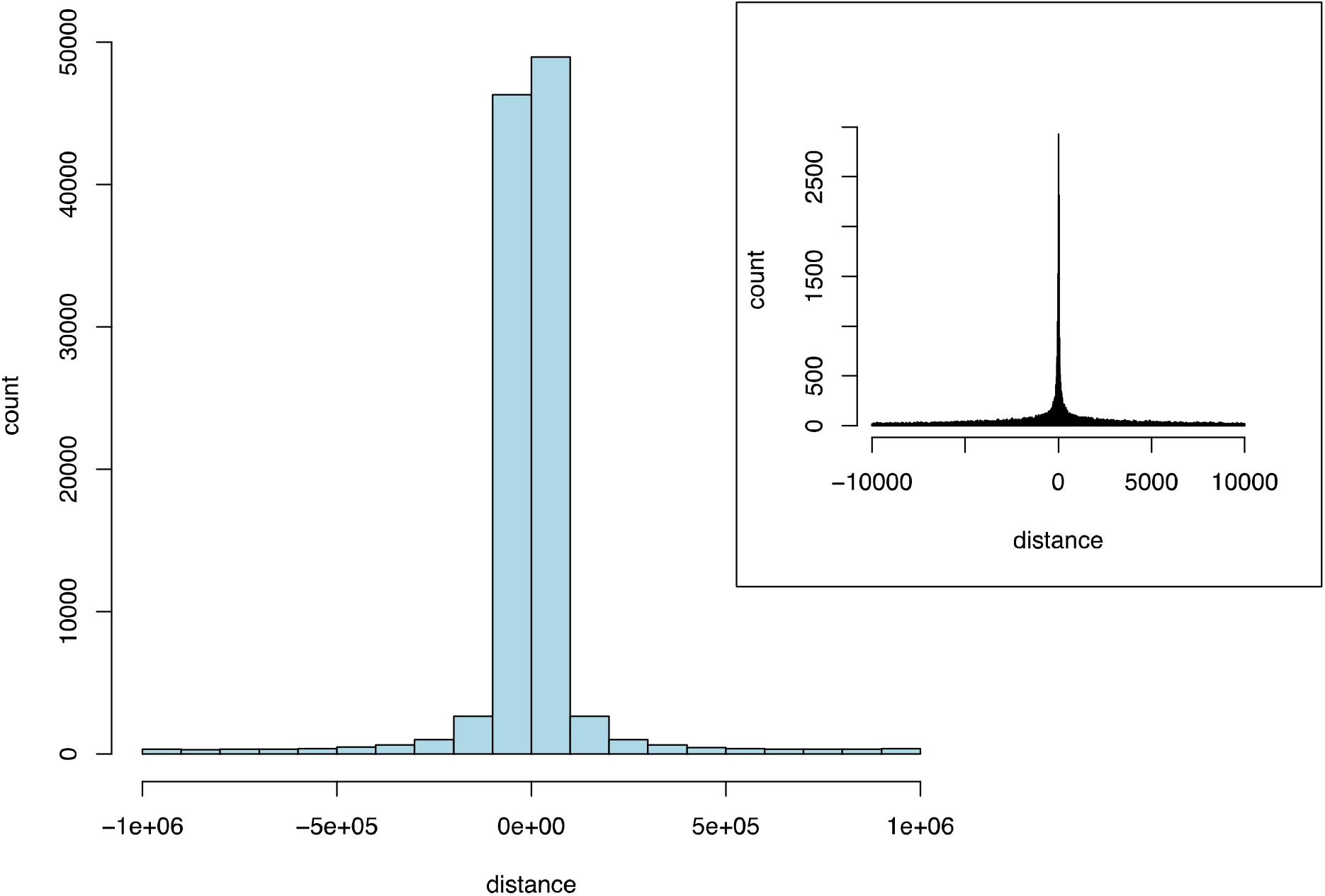
Distribution of distance between mSNPs and their target *cis*-mCpGs. Distance distribution is shown in base pairs (1Mb window). Inset: higher resolution illustration in ±10kb.

**Supplementary Figure 2.**
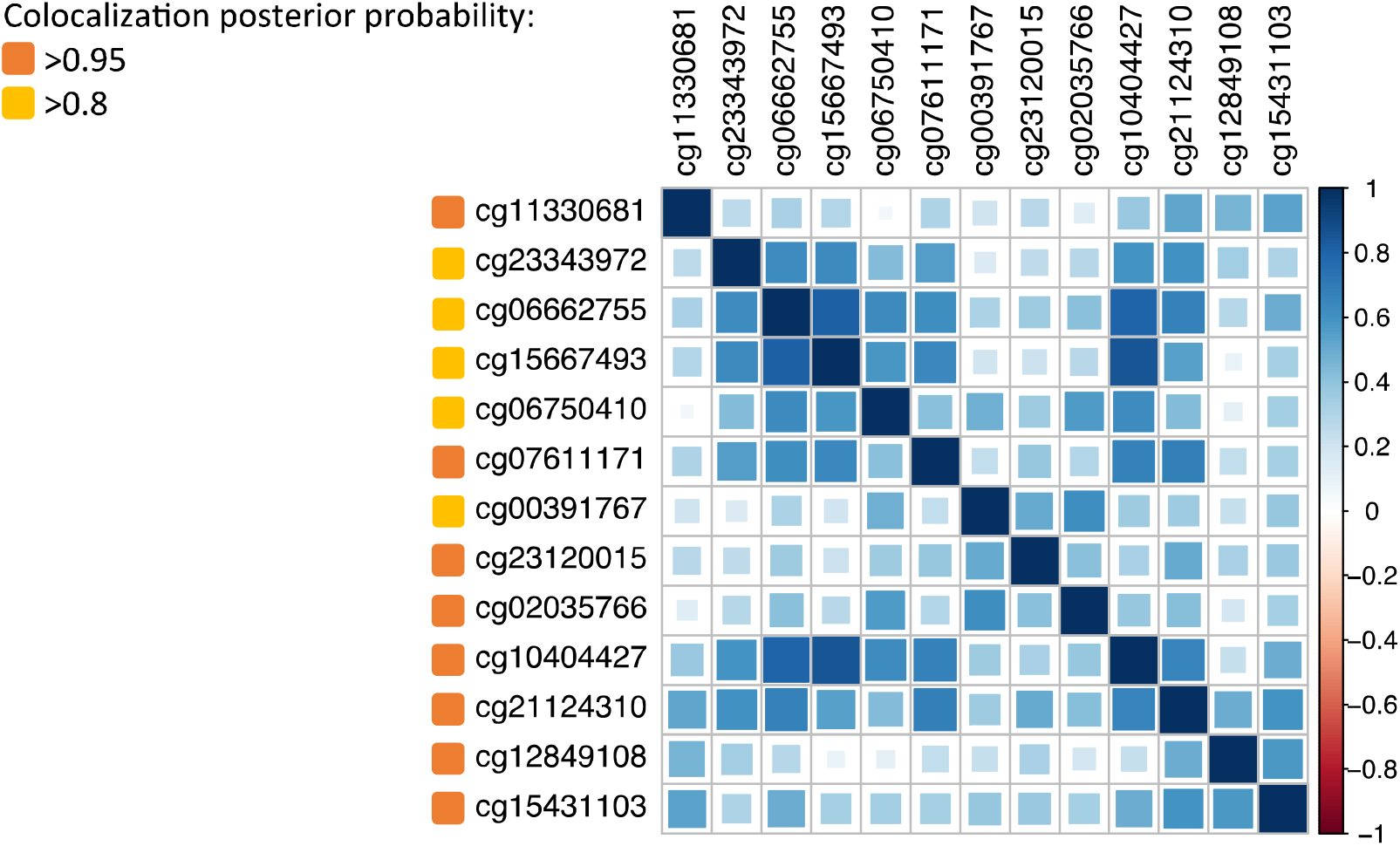
Correlation between methylation levels of CpGs affected by rs7731626 MS susceptibility locus. Methylation levels of all the colocalized mCpGs are positively correlated.

**Supplementary Figure 3.**
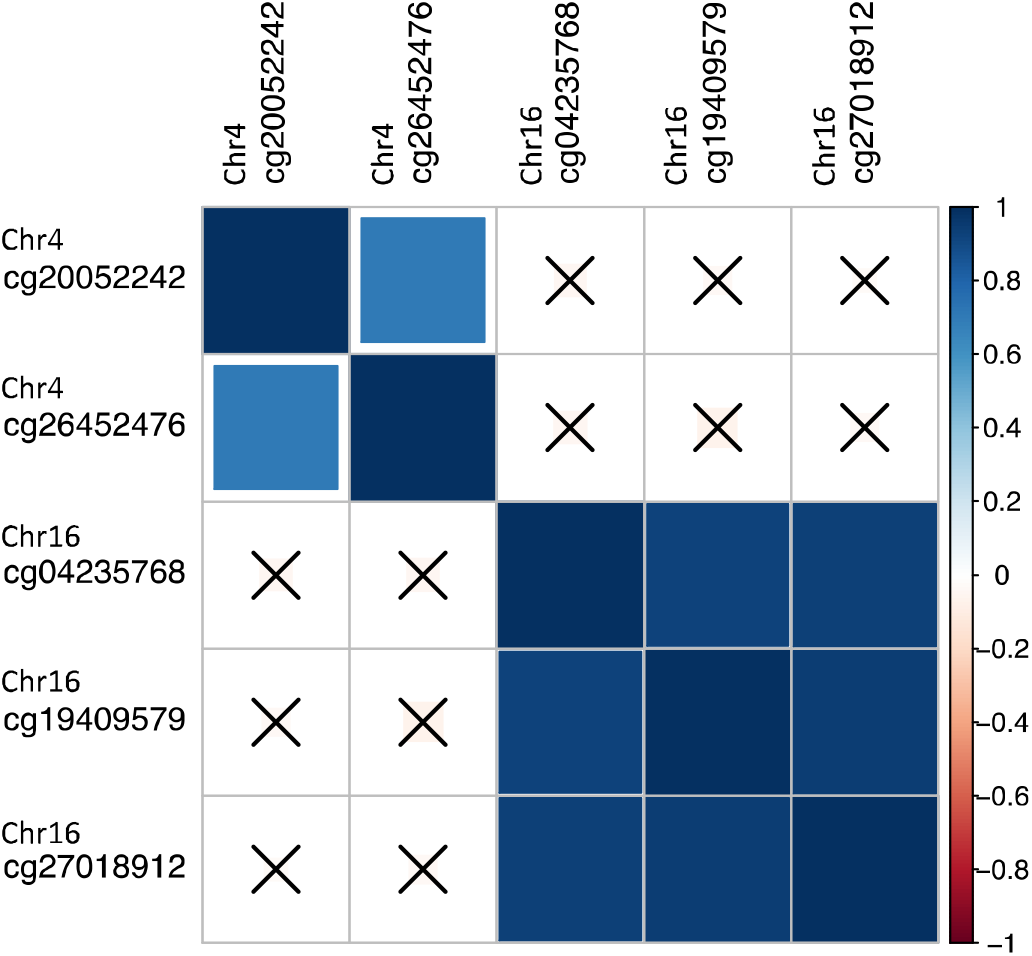
Correlation between *cis* and *trans* mCpGs affected by rs3809627 MS locus. No correlation is observed between methylation levels of the 2 *cis*-mCpGs on chromosome 16 and 3 *trans*-mCpGs on chromosome 4.

**Supplementary Figure 4.**
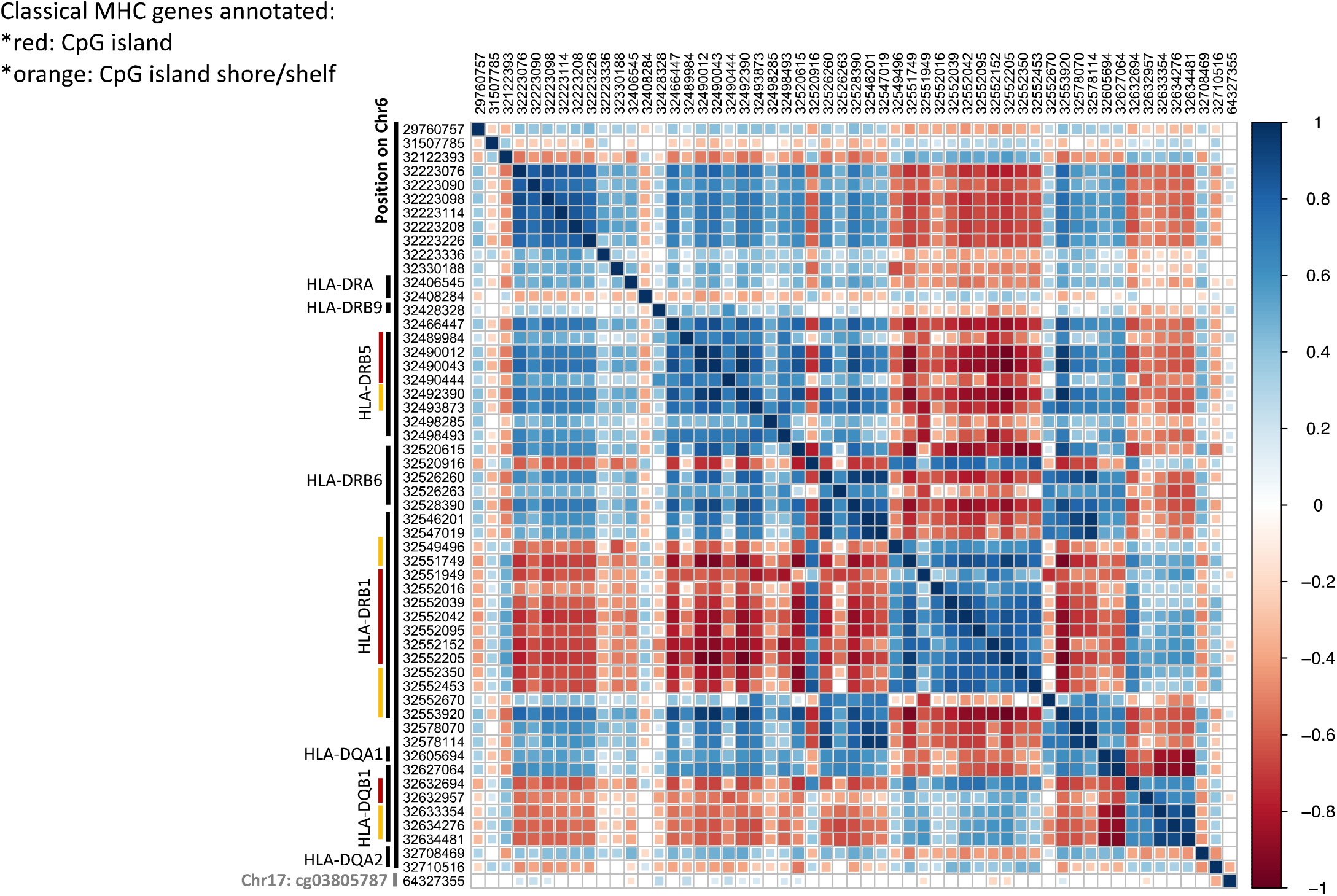
Correlation between methylation levels of CpGs affected by MS MHC polygenic score. Genomic coordinates are shown in GRCh37/hg19. All except one CpG are located in chromosome 6. The last CpG is located in chromosome 17.

**Supplementary Figure 5.**
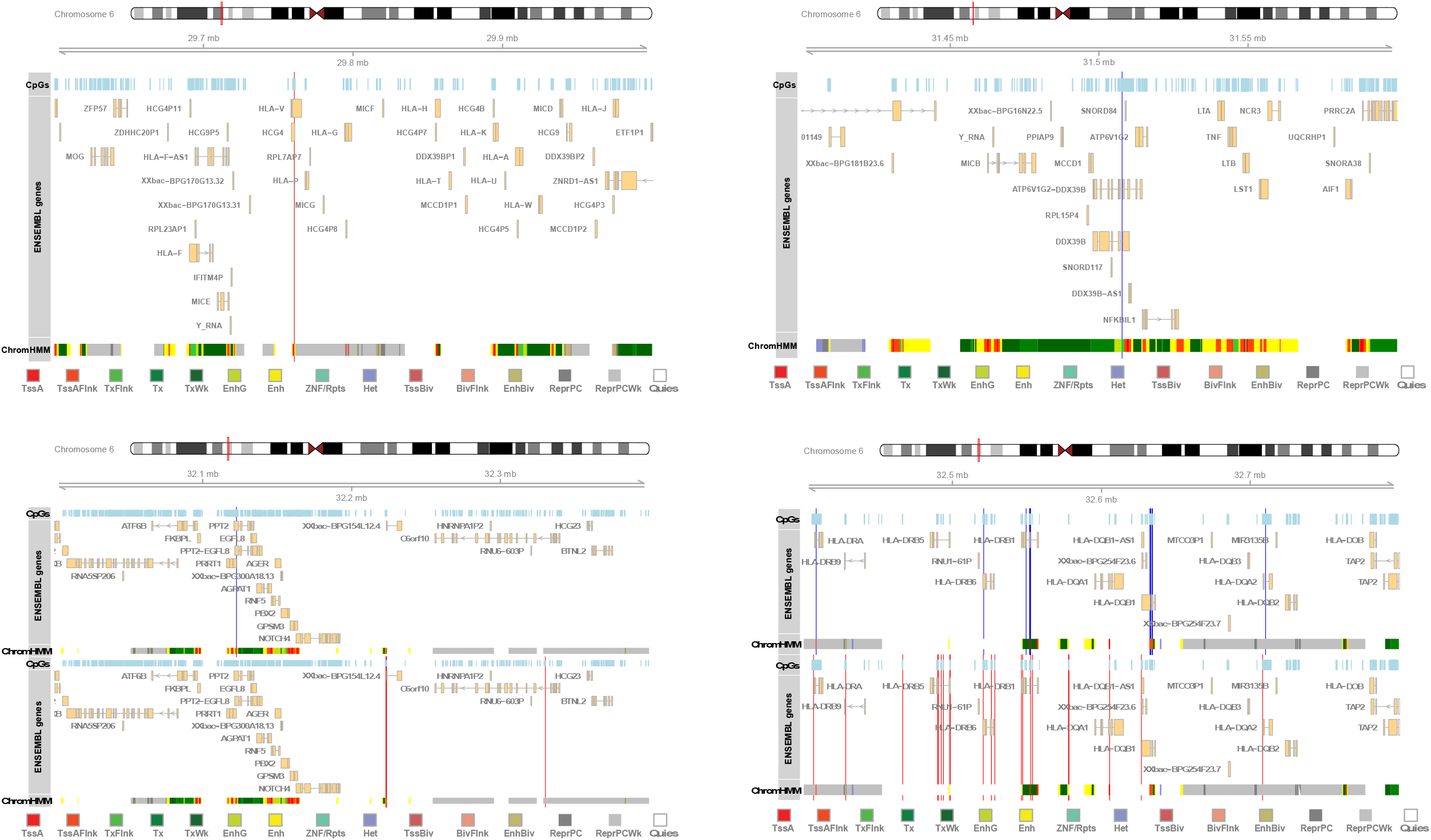
Genomic locations of MHC CpGs affected by MS MHC polygenic score in relation to nearby CpGs, genes, and modeled chromatin states. *Blue/red* vertical lines represent the locations for CpGs with negative/positive associations with the MHC polygenic score, respectively.

**Supplementary Figure 6.**
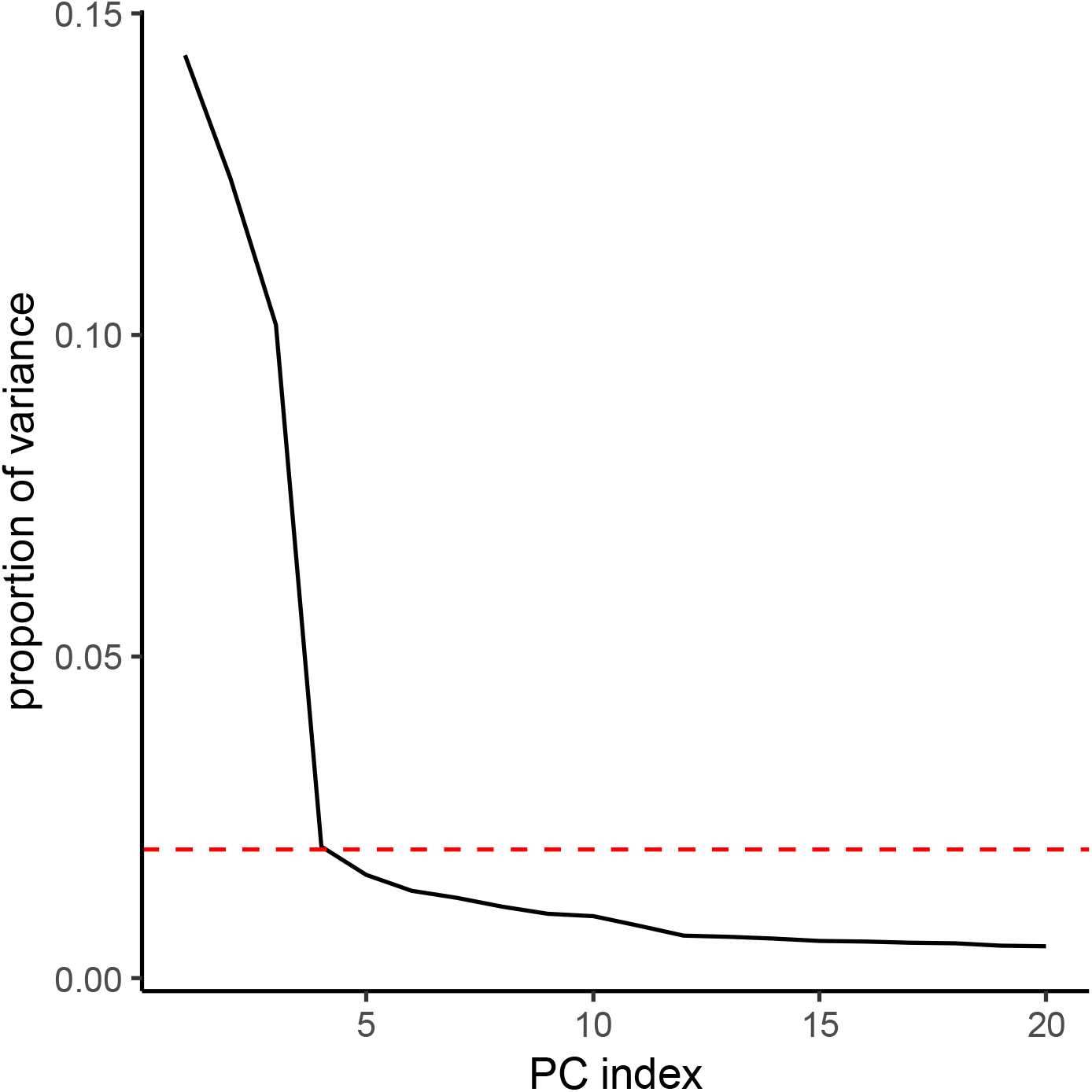
Variance explained by methylation PCs. Percentage of variance explained by the first 20 principal components of the methylation data is illustrated. *Red* dashed line marks 2%.

**Supplementary Table 1.** Subject demographics.

**Supplementary Table 2.** Genome-wide significant *cis*-mQTL effects.

**Supplementary Table 3.** Colocalized MS-mQTL effects.

**Supplementary Table 4.** *cis*-eQTL effects associated with colocalized MS-*cis*-mQTL effects.

**Supplementary Table 5.** CpGs affected by MS MHC polygenic score.

**Supplementary Table 6.** CpGs affected by MS total polygenic score.

**Supplementary Table 7.** MS susceptibility loci associated with cg16050799 methylation.

**Supplementary Table 8.** MS polygenic score calculation.

**Supplementary Table 9.** EpiTYPER analysis primers.

